# Upstream open reading frames may contain hundreds of novel human exons

**DOI:** 10.1101/2024.03.22.586333

**Authors:** Hyun Joo Ji, Steven L Salzberg

## Abstract

Several recent studies have presented evidence that the human gene catalogue should be expanded to include thousands of short open reading frames (ORFs) appearing upstream or downstream of existing protein-coding genes, each of which would comprise an additional bicistronic transcript in humans. Here we explore an alternative hypothesis that would explain the translational and evolutionary evidence for these upstream ORFs without the need to create novel genes or bicistronic transcripts. We examined 2,199 upstream ORFs that have been proposed as high-quality candidates for novel genes, to determine if they could instead represent protein-coding exons that can be added to existing genes. We checked for the conservation of these ORFs in four recently sequenced, high-quality human genomes, and found a large majority (87.8%) to be conserved in all four as expected. We then looked for splicing evidence that would connect each upstream ORF to the downstream protein-coding gene at the same locus, thus creating a novel splicing variant using the upstream ORF as its first exon. These protein coding exon candidates were further evaluated using protein structure predictions of the protein sequences that included the proposed new exons. We determined that 582 out of 2,199 upstream ORFs have strong evidence that they can form protein coding exons that are part of an existing gene, and that the resulting protein is predicted to have similar or better structural quality than the currently annotated isoform.

**Author Summary:** We analyzed over 2000 human sequences that have been proposed to represent novel protein-coding genes, and that reside just upstream of known genes. These “upstream ORFs” (uORFs) would represent a surprisingly large addition to the human gene catalogue, which after decades of refinement now contains just under 20,000 protein-coding genes. They would also create over 2000 new bicistronic genes, which number only 10 in current human annotation databases. We hypothesized that rather than novel genes, these sequences might instead represent novel exons that can be spliced into existing protein-coding genes, creating new isoforms of those genes. Using a combination of transcriptional evidence and computational predictions, we show that at least 582 of the previously-described uORFs can be used to create novel protein-coding exons, generating new transcripts and new protein isoforms, but not requiring the addition of entirely new genes to the human gene catalogue. We also demonstrate that the predicted three-dimensional structure of some of the new protein isoforms hints at new or improved functions for existing proteins.

## Introduction

Although the human protein-coding gene count has been converging on just under 20,000 genes in recent years (Amaral, Carbonell-Sala et al. 2023, Varabyou, Sommer et al. 2023), multiple recent studies have suggested the possible presence of thousands of additional short protein-coding genes (Ji, Song et al. 2015, Calviello, Mukherjee et al. 2016, Raj, Wang et al. 2016, van Heesch, Witte et al. 2019, Chen, Brunner et al. 2020, Gaertner, Van Heesch et al. 2020, Martinez, Chu et al. 2020, Mudge, Ruiz-Orera et al. 2022). Most of these proposed novel genes take the form of short open reading frames (ORFs) that occur just upstream or downstream of existing protein-coding genes, apparently on the same messenger RNA. Although thousands of these upstream ORFs (uORFs) have been hypothesized, only 25 have thus far been added to any of the major human annotation databases (Mudge, Ruiz-Orera et al. 2022).

One problem with the hypothesis that thousands of uORFs represent distinct proteins is that each of these would create a bicistronic transcript; i.e., a single transcript that encodes two distinct proteins. These are not unknown in the human genome, but they are very rare: as of now, only 10 bicistronic genomic loci have been annotated in the MANE (v1.3) annotation (Morales, Pujar et al. 2022), the current “gold standard” for human protein-coding gene annotation.

We therefore wished to examine a hypothesis that might explain many of the uORFs, but that would not require the addition of large numbers of novel genes and bicistronic transcripts to the human gene catalogue: what if these upstream ORFs represent protein-coding exons that have not previously been annotated? In this case, we would need to find evidence for splice sites that would link together each uORF with the known, downstream ORF at the same locus.

Previous work (Mudge, Ruiz-Orera et al. 2022) identified a set of uORFs that have at least two lines of evidence suggesting their protein-coding nature: (1) mutations within the uORFs that show a bias towards synonymous mutations; and (2) ribosome profiling data, in the form of Ribo-seq experiments (Brar and Weissman 2015), that suggest that the uORFs are being translated. Worth noting here is that Ribo-seq evidence cannot be used as direct evidence to prove the existence of a protein-coding sequence. A decrease in ribosome footprint density is typically interpreted as indicating a stop codon, but if a hypothesized uORF is part of an exon rather than a distinct gene, then a decrease might instead indicate a splice donor site; i.e., the end of the exon.

To search for novel exons that would explain uORFs, we considered several lines of evidence. First we confirmed that the novel exons were conserved in multiple distinct human genomes that have recently been sequenced. Second, we looked for direct evidence of splicing from the large-scale expression data generated by the GTEx project (Lonsdale, Thomas et al. 2013), and for computational evidence of splice donor sites within the uORFs that could be paired with previously-annotated acceptor sites. Either of these findings could link a uORF to a downstream protein-coding gene and create a novel isoform of that gene. Third, we evaluated the novel protein sequences created by each hypothesized new isoform, and we considered whether the folded protein structure, as predicted by ColabFold (Mirdita, Schütze et al. 2022), would be comparable to the structure of the canonical protein at that locus.

## Methods

We began our analysis with a large set of potentially novel human protein-coding regions collated from seven publications identifying ORFs from a variety of evidence sources, including Ribo-seq experiments and evolutionary conservation (Ji, Song et al. 2015, Calviello, Mukherjee et al. 2016, Raj, Wang et al. 2016, van Heesch, Witte et al. 2019, Chen, Brunner et al. 2020, Gaertner, Van Heesch et al. 2020, Martinez, Chu et al. 2020). These ORFs occur in regions previously thought to be untranslated, and in most cases appearing within the 5’ untranslated regions (UTRs) of protein-coding transcripts. A meta-analysis of these publications (Mudge, Ruiz-Orera et al. 2022) collected a total of 7,264 of these ORFs, including some contained entirely within known protein-coding regions and some that were in the 3’ UTR. 3,771 of the 7,264 ORFs were either entirely upstream or else partly upstream and partly overlapping with the known protein-coding region. For ease of discussion, we will refer to all of these as uORFs. We focused our analysis on a high-confidence subset of 2,199 uORFs that were reported by at least two publications.

### Preservation of uORFs in other human genomes

All uORFs considered here were originally identified on the reference human genome GRCh38. Because multiple additional human genomes have now been sequenced to high levels of completeness and accuracy, we reasoned that any novel protein-coding gene should be preserved in those other genomes as well. We evaluated the level of conservation by aligning the sequences from GRCh38 to each target genome, and then checking for any differences in the target genome. We also required that each uORF sequence be found in the 5’ UTR region of a transcript in the same gene locus on the target genome.

We aligned the uORFs to four different human genomes: a Puerto Rican individual (PR1, v3.0) (Zimin, Shumate et al. 2021), a Southern Han Chinese individual (Han1, v1.2) (Chao, Zimin et al. 2023), an Ashkenazi individual (Ash1, v2.2) (Shumate, Zimin et al. 2020), and the complete, gap-free CHM13 human genome (v2.0) (Nurk, Koren et al. 2022). We aligned each ORF sequence twice: first to the genome and then to the transcriptome (Figure 1). For both alignments, we used minimap2 (v2.26) (Li 2018) in its single-end short read (sr) mode. Each alignment result was then used to assign a conservation level on a scale from 0 to 7. If the genomic alignment and transcriptomic alignment results did not agree, the ORF with the maximum conservation level was used. Transcriptomic alignments helped to identify ORFs that spanned splice junctions, which could occur when the 5’ UTR region of a transcript contained more than one non-protein coding exon. See Supplementary methods for the specific parameters used for alignments.

**Figure 1.**
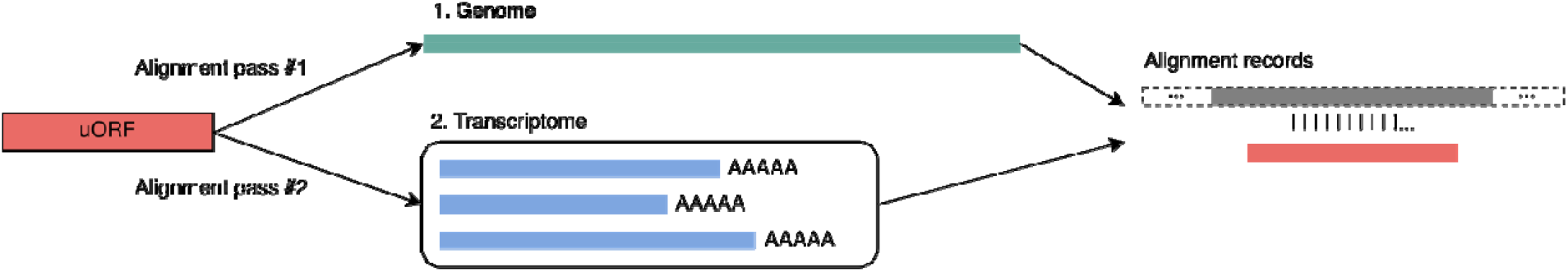
Each upstream ORF (uORF) was aligned to multiple human genomes, using both the genomic sequence and the annotated transcripts. The transcriptome alignment handled cases where a uORF spanned two exons in the 5’UTR of an annotated transcript.

As expected, most query ORFs yielded perfect nucleotide-level matches to the target genomes. For those that did not, we computed all translations and checked if any encoded a protein identical to the one encoded by the original query ORF. For each genome, there were handful of cases (35 on average) in which a query uORF had a perfect protein-level match without a sequence-level match. The resulting alignment coordinates were then annotated using the target genomes’ annotations. Genomic and transcriptomic alignments were processed slightly differently. For a genomic alignment, any overlapping gene and transcript features were collected. When an annotation was found, we checked to confirm that the uORF occurred in the 5’ UTR region of the annotated transcript and that the name of the downstream gene corresponded to the one described in the original source (Mudge, Ruiz-Orera et al. 2022).

To quantify conservation levels, we used three separate criteria with different weights, and the sum of the weights was used as a proxy for the level of conservation for uORFs (Table 1). All cases in which uORFs were assigned a score of 6, 4, 2, or 0 (i.e., lacking a gene locus match) were manually examined to ensure that they were not located in different genomic locations from what was annotated by Mudge et al. (Supplementary Methods). Any ORF scoring at least 4 was considered conserved, unless the sequence match occurred more than 1kbp away from the source gene. Using these criteria, 1931 out of the set of 2199 uORFs (87.8%) were conserved in all four target genomes (Supplementary Table S5).

**Table 1.** uORF conservation criteria and their weights. Each uORF was assigned a score between 0 and 7 and uORFs scoring > 3 were considered conserved in a genome.

|  |  |
| --- | --- |
| Sequence match | 4 |
| 5' UTR containment | 2 |
| Gene locus match | 1 |

### De novo construction of transcripts

All 2199 uORFs were evaluated for possible splicing evidence, even those excluded in the previous filtering step. Using either experimentally observed splice sites from GTex or predicted splice sites from the Splam program (Chao, Mao et al. 2023), we constructed putative novel coding sequences by concatenating the CDS from the uORF (up to the splice site) to the coding exons of the downstream gene, creating a novel *uORF-connected transcript* (**Figure 2**). Each uORF-connected transcript was associated with a reference transcript from the GENCODE (v39) annotation and with the protein sequence annotated for that transcript.

**Figure 2.**
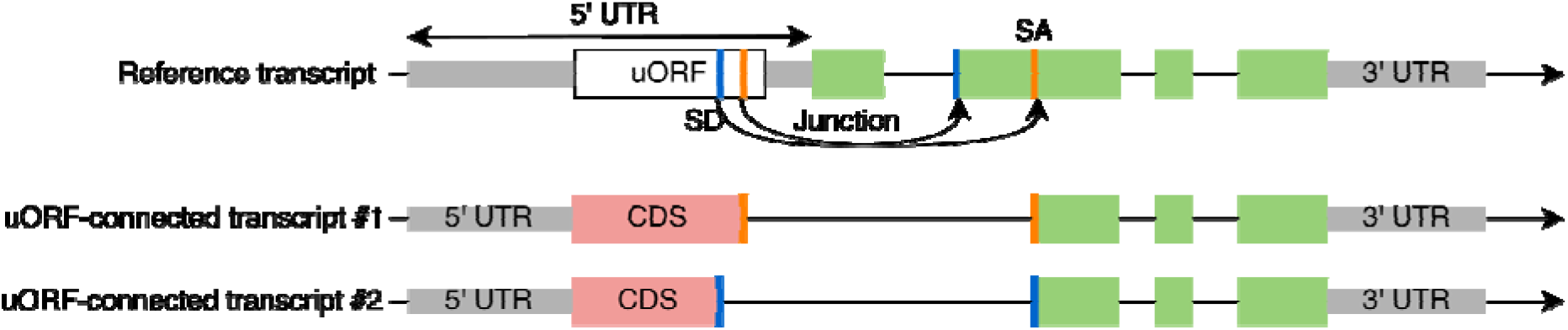
Construction of uORF-connected transcripts from a uORF and a downstream protein-coding transcript. The original protein-coding sequence is shown in green rectangles. For uORF-connected transcript #1, a splice junction (red bars) found in the GTEx collection of RNA-seq data is used to link the uORF to the second exon of the downstream transcript. For uORF-connected transcript #2, a splice donor (SD) site predicted by Splam (blue bars) is paired with an annotated splice acceptor (SA) site in a MANE transcript. The novel protein sequences are shown in red rectangles.

### Incorporation of experimental splicing evidence and predicted splice junctions

We collected our experimentally identified splice junctions from a large set of assembled transcripts created as part of the initial construction of the CHESS human annotation (Varabyou, Sommer et al. 2023), which is based on assemblies of 9,814 RNA-seq samples collected by the GTEx project (Lonsdale, Thomas et al. 2013). Note that most of these splice sites do not appear in the final CHESS catalogue. We merged and summarized the assembled transcripts using TieBrush (Varabyou, Pertea et al. 2021), which also captured the read coverage for each junction.

If a splice junction’s donor site occurred within a uORF and its acceptor site was contained in the downstream ORF at the same gene locus, we then attempted to connect the upstream and downstream ORFs to yield a novel ORF (**Figure 2**). We discarded any such construct if (1) it contained a premature stop codon, (2) its length was not a multiple of three, or (3 it encoded a CDS whose length was < 90% of the length of the reference transcript. This filtering ensured that each putative novel isoform would produce a valid protein sequence of similar length to the reference protein.

We then considered additional splice sites (i.e., those not seen in GTEx) if they were given high scores by Splam, a highly accurate splice site predictor (Chao, Mao et al. 2023). We created these potential splice sites by first scanning the uORF regions to detect potential splice donors (GT dinucleotides), and then pairing them with splice acceptors coming from annotated MANE transcripts. Each junction was then scored by Splam, and we retained those with an average score for the donor and acceptor of at least 0.9. We then applied the same filtering steps as described above, removing ORFs if they would not generate a protein of similar length to the reference.

There were total of 166,465,454 possible junctions in the GTEx-supported set and 4,928 additional junctions in the SPLAM-predicted set. We created 2,977 candidate transcripts using TieBrush junctions and an additional 1,903 using the Splam-predicted spliced junctions (**Figure 3**). In total, we constructed 4,880 uORF-connected transcripts from 1,102 uORFs, provided in Supplementary Table S2.

**Figure 3.**
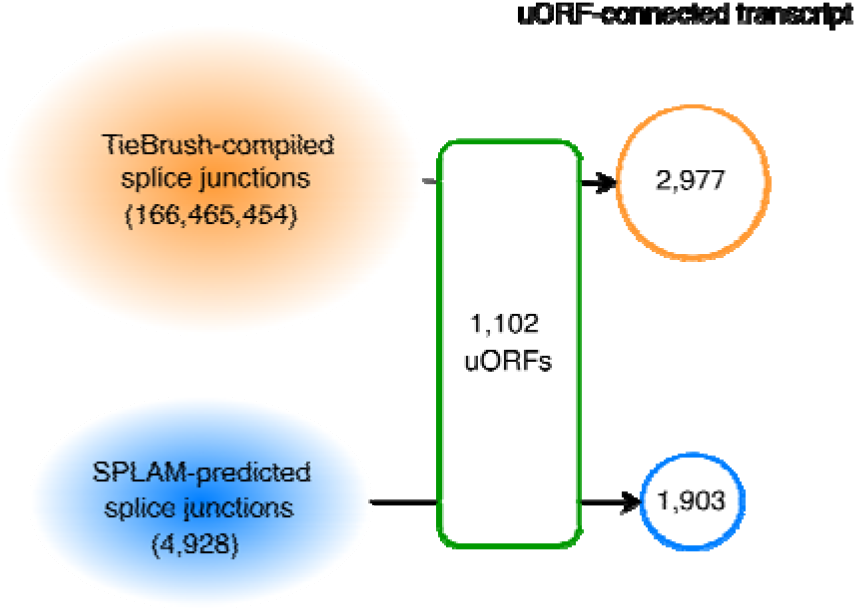
Splice junctions seen in ∼10,000 GTEx RNA-seq and Splam predictions yielded a total of 4,880 uORF-connected transcripts from 1,102 uORFs. 2,977 were supported by GTEx data and 1,903 had Splam support but not GTEx evidence.

In a handful of cases (26), uORF-connected transcripts encoded proteins with a non-AUG start codon, or more specifically, isoleucine (I) instead of methionine (M). These transcripts were included in our analysis but are separately annotated, in Supplementary Table S7. We also noted that a very small number of novel transcripts encoded proteins with exactly the same amino acid sequence as their references (due to duplications between the uORF and the downstream ORF), and these are separately shown in Supplementary Table S8.

### Folding transcripts

We used ColabFold, a tool that accelerates AlphaFold2’s protein structure predictions (Jumper, Evans et al. 2021, Mirdita, Schütze et al. 2022), to predict the structure of all novel proteins encoded by uORF-connected transcripts. We used the pLDDT score, AlphaFold2’s per-residue confidence metric, as a proxy for the quality of each folded structure. As described in the AlphaFold2 publication (Jumper, Evans et al. 2021), a protein with pLDDT >70 is considered to have high confidence and stability. See Supplementary methods for the specific parameters used for folding proteins using ColabFold.

If a uORF-connected transcript encoded a protein will a similar pLDDT score to the one encoded by its reference, then we considered it more likely that the novel protein was a valid functional isoform. In contrast, if the novel protein had a lower pLDDT score, then we eliminated it from further consideration (although some of these may nonetheless be valid). More precisely, a uORF-connected transcript that encoded a protein that achieved either an increase (>1) or no change (±1) in average pLDDT was retained for further investigation. Occasionally, a uORF-connected transcript was compared to multiple reference proteins when the locus contained multiple alternative proteins. In that case, we required that the uORF-connected transcript have a similar or better pLDDT than at least one of the references.

## Results

### Most uORFs are conserved in other human genomes

As described in Methods, we checked all uORFs to determine if they were conserved in four different human genomes: PR1, Han1, Ash1, and CHM13. Because humans are very closely related, we only considered a uORF to be conserved in another human if it had a complete sequence-level match; i.e., full-length and 100% identical. Additional criteria such as the uORF’s containment in the 5’ UTR region or consistency of the gene locus was used to further classify uORFs into different levels of conservation (Table 2 and Methods).

**Table 2.** Number of uORFs in GRCh38 that were conserved in each of four other human genomes, at varying levels of conservation. Level 7 indicates a perfect match contained in the 5’ UTR region of a protein-coding transcript at the correct gene locus. Values in parentheses are the number of cases in which the best sequence match occurred at a different genomic location (>1kbp away), not the one previously annotated.

| Conservation score |  | Han1 | CHM13 | Ash1 | PR1 |
| --- | --- | --- | --- | --- | --- |
| Conserved | 7 | 1,931 | 1,943 | 2,078 | 2,065 |
|  | 6 | 14 | 15 | 1 | 2 |
|  | 5 | 66 | 68 | 6 | 4 |
|  | 4 | 25 (5) | 27 (5) | 5 (0) | 2 (0) |
|  | Subtotal | 2,036 (5) | 2,053 (5) | 2,090 | 2,073 |
| Not conserved | 3 | 146 | 131 | 108 | 125 |
|  | 2 | 2 | 1 | 0 | 0 |
|  | 1 | 4 | 5 | 0 | 0 |
|  | 0 | 6 | 4 | 1 | 1 |
|  | Subtotal | 158 | 141 | 109 | 126 |

A large majority of the uORFs, 1931 out of 2199 (87.8%), were conserved in all four individual human genomes as well as GRCh38 (Figure 4). Most of the remaining uORFs (213/268) were conserved in at least two of the four genomes, with only a handful conserved in exactly one genome. Interestingly, there were 43 ORFs shared only between Ash1 and PR1, 36 conserved in all genomes except PR1, and 30 conserved in all genomes except Han1. The conservation levels for each of the 2,199 uORFs can be found in Supplementary Table S5.

**Figure 4.**
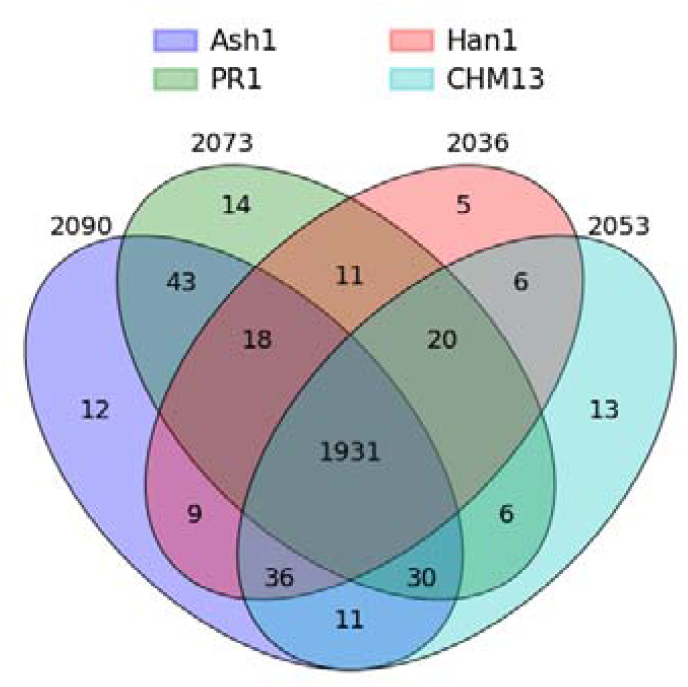
Conserved uORFs shared between GRCh38 and all subsets of four different genomes. The innermost region shows that there were 1,931 uORFs conserved in all five genomes.

### uORFs give rise to novel protein coding exons

Our main hypothesis is that some uORFs identified within known protein-coding transcripts might instead be explained as novel protein-coding exons that represent isoforms of the existing gene, which a priori seemed more likely than an entirely new protein-coding gene sharing the same transcript. As described in Methods, we constructed novel exons using splice junctions supported by either experimental (RNA-seq) or computational evidence, and retained transcripts only if they encoded a protein at least 90% as long as the reference.

Out of 2,977 proteins encoded by uORF-connected transcripts with GTEx support, 1,035 had a pLDDT score that was the same or higher than their reference proteins. The RNA-seq read coverage on the novel junctions in these 1,035 uORF-connected transcripts was higher on average (143) than the mean coverage for all junctions used to construct uORF-connected transcripts (117). These transcripts corresponded to 352 distinct uORFs from the original set.

Although many fewer uORF-connected transcripts (1,903) were constructed using Splam-predicted junctions, 757 of these encoded proteins with an equal or higher pLDDT score than their references. These Splam-supported transcripts corresponded to 314 distinct uORFs, of which 84 were also used to construct GTEx-supported transcripts. Splam was used to score all junctions not included in the splice junction set extracted from GTEx samples, so this overlap represents cases where a Splam-supported junction and a GTEx-supported junction link the same uORF to its downstream protein. The union of these two sets contained 582 unique uORFs from the original set of 2,199 uORFs proposed by Mudge et al. (Mudge, Ruiz-Orera et al. 2022).

In total, we constructed 4,880 uORF-connected transcripts encoding distinct proteins with either GTEx or Splam support. 1,792 of these encoded proteins with a pLDDT score that was the same or higher than their references, and we further investigated 529 of the 1,792 instances with strictly higher average pLDDT scores. The structures for these 529 proteins were manually compared their references and categorized into five groups: end truncation, alpha helix elongation, straightening, tightening, and structural addition. Some proteins displayed more than one of these changes. The most common type of change was end truncation (189), followed by alpha helix deletion / truncation (132), structure addition (41), alpha helix elongation (26), alpha helix straightening (7), and tightening (6). For cases of alpha helix deletion / truncation that resulted in an increase in average pLDDT, we observed that the structure deletions were often accompanied by removal of long, unstructured coils, as illustrated for the OPA1gene locus in Supplementary Figure S1B.

Figure 5 illustrates several of the structural improvements we objected. In Figure 5A-B, an alpha helix was lengthened in a uORF-connected isoform of SLC28A1, which led to an increase of 2.96 in the pLDDT score. In Figure 5C-D, we illustrate a “straightening” event, where two alpha helix structures in the native isoform of TRAK2 were interrupted by a kink and a short coil region. In the uORF-connected isoform, these were connected and formed a single helix, which yielded an increase of 3.41 in pLDDT. Figure 5E-F illustrates a cases in which the structural components of a protein were brought closer together, or “tightened.” Tightening is particularly interesting for proteins with binding domains, as it might yield an increased binding efficacy for a protein to its target. The ORF-connected transcript at the TBRG4 gene locus shown in Figure 5F encodes a protein predicted to have a tighter conformation, leading to an increase of 4.29 in average pLDDT. The TBRG4 protein, also known as FASTKD4, contains an RNA-binding domain (RAP), whose efficacy may be influenced by how tightly the protein is packed (Simarro, Gimenez-Cassina et al. 2010, Yeung, Das et al. 2011).

**Figure 5.**
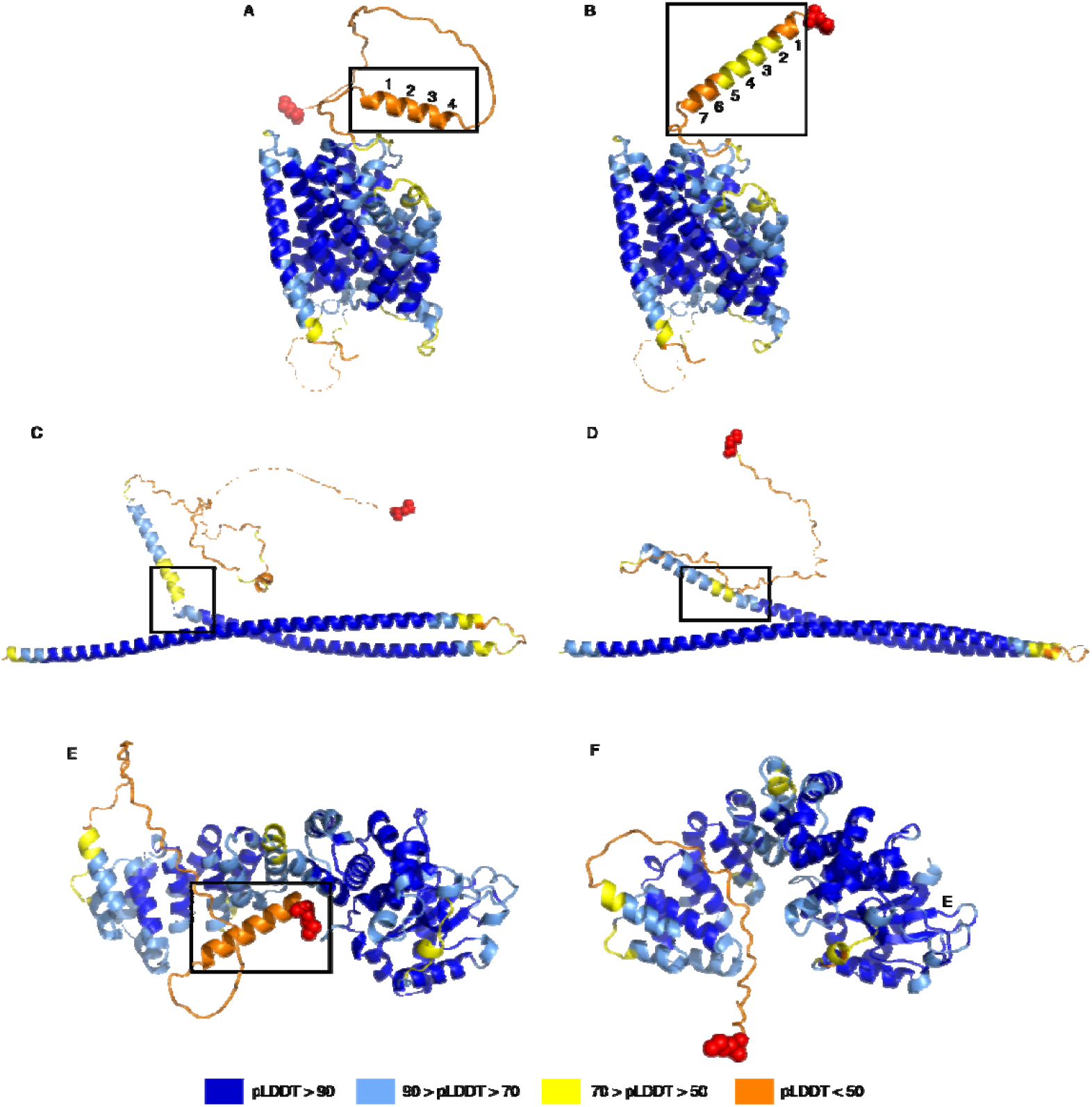
Examples of structure changes in novel protein variants identified in this study. (A) and (B): alpha helix elongation at the SLC28A1 gene locus, where (A) shows the reference protein, ENST00000398637.10, and (B) shows the novel isoform, uorft_2119. The average pLDDT increase from A to B was 2.96. (C) and (D): straightening at the TRAK2 gene locus, where (C) shows the native protein, ENST00000430254.1, and (D) shows the novel isoform, uorft_441. The average pLDDT increase from C to D was 3.41. (E) and (F): tightening of a structure of TBRG4, where (E) shows the known protein, ENST00000395655.8, and (F) shows the novel isoform, uorft_1435. The main structural changes are highlighted by black boxes for each pair of structures. Red spheres represent the N-terminus of each protein. The average pLDDT increase from E to F was 4.29.

The most common type of change we observed was end truncation, where an unstructured (coil) region at the N-terminus of a protein was removed, resulting in a higher average pLDDT score. For instance, the isoform defined by a uORF-connected transcript at the ZDHHC5 gene locus has a much shorter coil compared to the canonical protein at that locus, leading to an increase of 4.5 in average pLDDT score (Supplementary Figure S1A).

Finally, structure addition refers to cases in which one or more alpha helices or beta sheets were added to the reference protein. Adding well-folded structures to a protein generally leads to an increase in its pLDDT score. For example, a uORF-connected transcript at the OXNAD1 gene locus adds an alpha helix that yields an increase of 3.63 in average pLDDT (Supplementary Figure S1C). Similarly, another uORF-connected transcript at a different gene locus, ZNF32, adds a series of four beta sheets near the N-terminus of the reference protein, yielding an increase of 3.28 in average pLDDT (Supplementary Figure S1D).

## Discussion

We evaluated 2,199 upstream ORFs that were previously proposed as novel human proteins, all of them on existing protein-coding transcripts. If these uORFs turn out to be genuinely novel genes, they will also create bicistronic transcripts, representing a huge increase in the number of such transcripts in the human genome, which currently number only 10. Our analysis suggests that at least 582 of these uORFs (26% of the total) could instead be used to form novel protein-coding exons that can be connected to exons from existing genes to form new protein variants. These new isoforms can be added to the human genome annotation without creating entirely new genes or bicistronic transcripts. Our process incorporated multiple lines of evidence including RNA sequencing support, inter-individual sequence conservation, computational splice site predictions, and protein structure prediction. We should emphasize that our process was conservative, and the remaining 1,617 uORFs might also represent protein-coding exons. For example, we required that the predicted structure of each novel isoform have either the same or higher average pLDDT than the reference, but isoforms with a lower pLDDT might nonetheless be valid. Further research is necessary to determine if these additional uORFs are novel proteins or instead might be merged with existing genes.

## Data Availability

The code used to produce the results and conduct analyses presented in this manuscript is publicly available at https://github.com/haydenji0731/uORF_explorer under the GPLv3 license, which is also archived on Zenodo at link https://zenodo.org/doi/10.5281/zenodo.10854712. The analysis results, including GTF files containing the annotation of uORF-connected transcripts and the PDB files for the protein structures included as part of this manuscript, are also available at the same GitHub address. Rest of the relevant data are within the paper and its Supporting Information files.

## Acknowledgements

We would like to thank Ales Varabyou for all the useful discussions on uORFs and generating the TieBrush output on the GTEx RNA-seq data. This work was supported in part by the US National Institutes of Health under grants R01-HG006677, R01-MH123567, and R35-GM130151.

## Contributions

HJJ and SLS designed the study. HJJ wrote the code used for analyzing uORFs, including protein structure predictions. HJJ and SLS evaluated analysis results and wrote manuscript drafts. All authors read and approved the final manuscript.

