## Supplemental Figures & Methods for "Upstream open reading frames may contain hundreds of novel human exons"

### Supplementary Figures

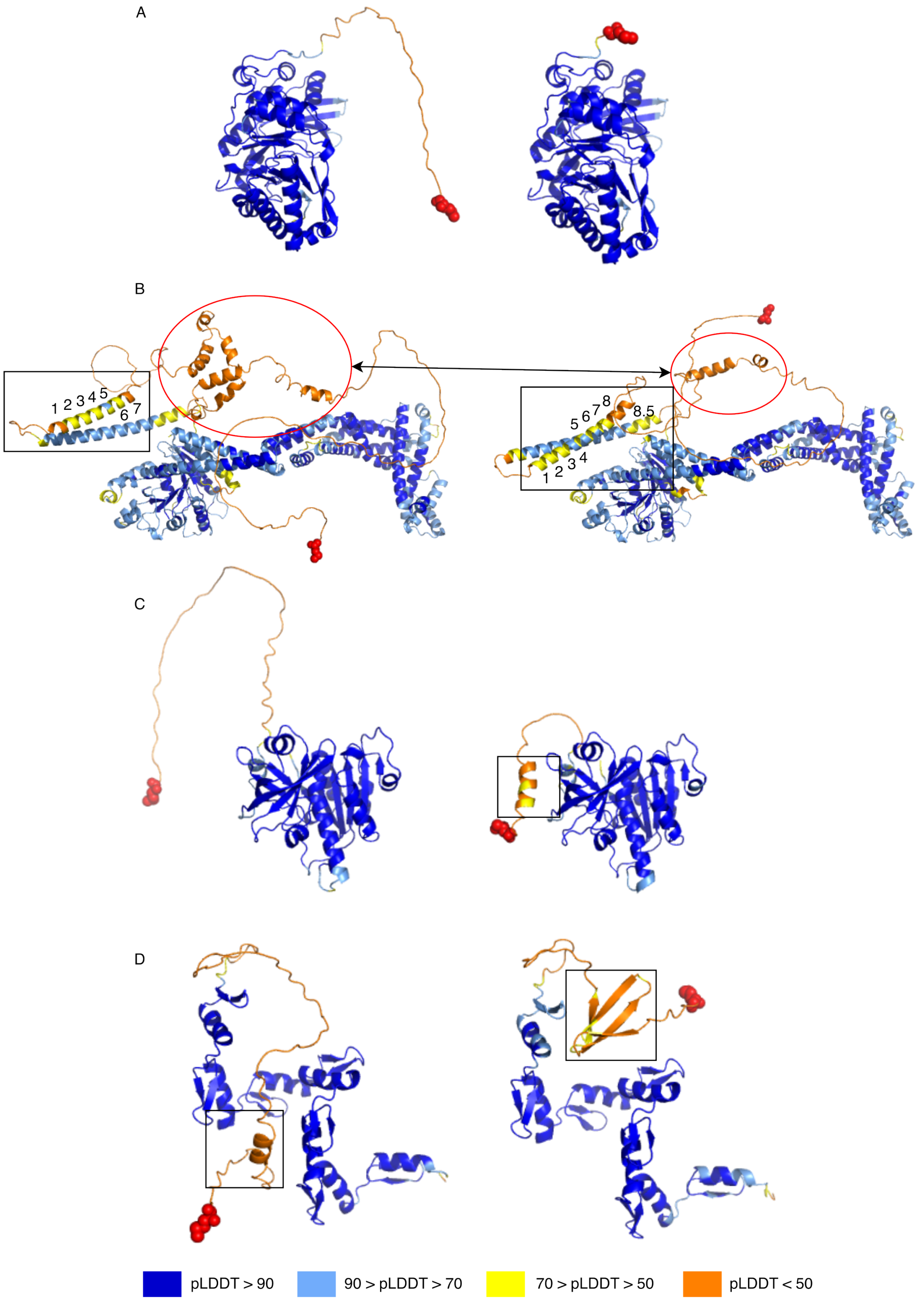


Supplementary Figure S1. Additional examples of structure changes in novel protein variants identified in this study. (A): Coil truncation at the ZDHHC5 gene locus. The reference protein (ENST00000323578.13) is on the left and novel isoform (uorft_2063) is on the right. The pLDDT increase was 4.5 (B): Alpha helix deletion at the OPA1 gene locus. The reference protein (ENST00000361510.8) is on the left and novel isoform (uorft_760) is on the right. The pLDDT increase was 3.23. (C): Alpha helix addition at the OXNAD1 gene locus. The reference protein (ENST00000285083.10) is on the left and novel isoform (uorft_704) is on the right. The pLDDT increase was 3.63. (D): Alpha helix was replaced by beta sheets at the ZNF32 gene locus. . The reference protein (ENST00000374433.7) is on the left and novel isoform (uorft_1781) is on the right. The pLDDT increase was 3.28. The main structural changes are highlighted by black boxes for each pair of structures. Red spheres represent the N-terminus of each protein.

### Supplementary Methods

#### uORF sequence alignment with minimap2

Each uORF sequence was treated as a short single-end read. Hence, we used minimap2's sr preset to align uORF sequences to either the genome sequence or annotated transcript sequences. When using mappy, the python wrapper for minimap2 is used, this looks like:

```
mappy.Aligner(fn_idx_in=genome.fa, fn_idx_out=index_file, preset=”sr”)
mappy.Aligner(fn_idx_in=transcripts.fa, fn_idx_out=index_file, preset=”sr”)
```

When using the minimap2 binary, this becomes:

```
minimap2 -ax sr genome.fa uORF.fa
minimap2 -ax sr transcripts.fa uORF.fa
```

#### Protein structure prediction with ColabFold.

To fold the proteins defined by uORF-connected transcripts, we ran the local version of the ColabFold available at https://github.com/YoshitakaMo/localcolabfold. We used the default parameters suggested by the ColabFold authors for monomer predictions and have additionally set the random seed to 0 for all predictions to increase the reproducibility of our predictions. Predictions were made using NVIDIA A100 GPU.

ColabFold and AlphaFold2 predictions can be time consuming for long protein sequences of length > 1,000 . For practical reasons, we skipped proteins of length > 2,000 amino acids when folding uORF-connected transcripts with Splam support.

#### Manual curation of uORF mapping positions.

When checking for the conservation of a uORF sequence in a genome, we looked for a sequence-level match and if there's match, then its location was examined. If the uORF sequence is mapped to a completely different genomic location (i.e., far from the gene locus that Mudge et al. annotated), then it wasn't considered conserved. The boundaries for a gene locus can vary depending on the annotation, so we further analyzed uORFs that were flagged at being outside of the previously annotated gene locus, or source gene for simplicity. More specifically, we computed the distance between where these uORFs were mapped and their source genes. If this distance was small, then we had more reasons to be believe that corresponding uORF is conserved.

Distances were defined as the number of bases between uORF alignment end position and the source gene locus start position (see Supplementary Table S4). Surprisingly, some genes were missing from the annotation, which had caused some uORFs to be flagged because there's no gene locus to match for. We observed that most missing genes had the name prefix “AC” or “AL”. All of them were overlapping with a more canonical gene, indicating that a uORF on one of these loci is in fact associated with multiple genes. Hence, we obtained the genomic coordinates for these missing genes by taking the genomic coordinates of the overlapping gene locus or aligning their transcript sequences to the target genome using minimap2. For few instances, the distances were either negative (i.e., the gene and uORF are overlapping) or marked as “c” (i.e., uORF is contained in the gene locus). All uORF matches that occur less than 1000 bp away from their source gene loci were counted as conserved.
